# MRI mapping of hemodynamics in the human spinal cord

**DOI:** 10.1101/2024.02.22.581606

**Authors:** Kimberly J. Hemmerling, Mark A. Hoggarth, Milap S. Sandhu, Todd B. Parrish, Molly G. Bright

## Abstract

Impaired spinal cord vascular function contributes to numerous neurological pathologies, making it important to be able to noninvasively characterize these changes. Here, we propose a functional magnetic resonance imaging (fMRI)-based method to map spinal cord vascular reactivity (SCVR). We used a hypercapnic breath-holding task to evoke a systemic vasodilatory response during concurrent blood oxygenation level-dependent fMRI. SCVR amplitude and hemodynamic delay were mapped at the group level as proof-of-concept of the approach, and in two highly-sampled participants to probe feasibility/stability of individual SCVR mapping. Across the group and individuals, a strong ventral SCVR amplitude was initially observed without accounting for local regional variation in the timing of the vasodilatory response. Shifted breathing traces were used to account for temporal differences in the vasodilatory response across the cord, producing maps of SCVR delay. These maps demonstrate distinct gray matter regions concordant with territories of arterial supply. The SCVR fMRI methods described here enable robust mapping of spatiotemporal hemodynamic properties of the human spinal cord. This noninvasive approach has the potential to provide early insight into pathology-driven vascular changes in the cord, which may precede and predict future irreversible tissue damage and guide the treatment of several neurological pathologies involving the spine.

## Introduction

Impaired vascular function is a major contributor to multiple neurological pathologies involving the spinal cord, including traumatic spinal cord injury ^1^ and degenerative cervical myelopathy ^2^. It is therefore of great importance, for both basic research and clinical translation, to characterize these changes in blood supply to the spinal cord tissue. However, existing methods to noninvasively characterize vascular function in the human brain have not yet been successfully translated to the cord. In general, physiological and functional magnetic resonance imaging (fMRI) of the spinal cord has lagged the development of such methods in the brain. This has largely been due to the challenges typically associated with these methods, including the small size of the spinal cord, nearby sources of physiological noise, magnetic field inhomogeneities, low signal to noise ratio (SNR), and lower widespread availability of specialized radiofrequency coils ^3,4^. More recently, technological and methodological developments have enabled more robust fMRI studies of the spinal cord, including mapping of sensory, motor, and resting-state neural activation patterns ^5^.

Despite these advances, MRI techniques for characterizing vascular properties of the spinal cord have remained somewhat limited. Dynamic susceptibility contrast (DSC) MRI, following injection of a gadolinium-based contrast agent, presents the gold standard for quantifying tissue perfusion and blood volume in the brain. Existing studies of vascular changes in the spinal cord of people with degenerative cervical myelopathy use this technique, showing sensitivity to the ischemic effects of cord compression ^6,7^. However, due to the low SNR of DSC data in the cord, these works report a single non-quantitative value for the entire cord region, rather than quantifying and mapping absolute vascular metrics. Repeated scans may boost data quality, but they are not readily feasible in DSC due to the long wash-out time of the contrast agent and the growing safety concerns over repeated exposure to gadolinium-based contrast agents ^8^. Dynamic contrast enhanced (DCE) MRI measures the contrast agent accumulation ^9^; although more commonly used to look at tumors in the spine, preliminary work demonstrates the potential of DCE to evaluate spinal cord perfusion after compression, but the technique is still in early development ^10^.

A contrast-free alternative, arterial spin labeling (ASL) MRI, shows excellent promise in the spinal cord injured rat ^11,12^, and in the human spinal cord in early conference proceedings ^13,14^. However, despite rapid growth of ASL for perfusion mapping of the brain, the unique challenges of spinal cord ASL have continued to hinder progress (e.g., low SNR, variable vascular anatomy for labelling and delay modeling, B_0_ inhomogeneity influencing labeling efficiency) ^11^. Recent exciting advances in pre-clinical studies are renewing interest and enthusiasm for spinal cord ASL in humans, but substantial methodological development is still needed before this technique can reliably shed light on human spinal cord pathophysiology ^11^. Another current research area is high-field intra-voxel incoherent motion (IVIM) MRI, which has shown promising maps of healthy human spinal cord perfusion, albeit in a single slice ^15^.

Here, we propose an alternative technique to map vascular function throughout the human spinal cord using blood oxygenation level-dependent (BOLD) fMRI, a well-established neuroimaging technique that is sensitive to changes in regional blood flow, and which has been successfully adapted and optimized for spinal cord neural activation mapping in recent years ^5^. Specifically, we use fMRI to map spinal cord vascular reactivity (SCVR), the cord’s blood flow responsiveness to a systemic vasodilatory stimulus. In the brain, vascular reactivity and hemodynamic delay measured with BOLD fMRI have been established as robust metrics of cerebrovascular function ^16,17^. However, techniques for adapting these cerebrovascular reactivity methods to the spinal cord are not yet well established. Two prior spinal cord fMRI studies have used gas inhalation challenges to modulate the vascular system; however, these are smaller studies (<10 participants) without more modern acquisition and modeling innovations, and do not map the distribution of vascular reactivity and delay ^18,19^.

In this study, we use advanced spinal cord fMRI methods to measure the local vasodilatory response to hypercapnia and map SCVR across the cervical spinal cord in healthy adults. We evoke systemic hypercapnia (elevated CO_2_) with simple breath-holding tasks (see reviews on breath-hold cerebrovascular reactivity ^16,17^), precluding the need for gas delivery systems. By recording end-tidal CO_2_ (P_ET_CO_2_) levels during scanning, we form a subject- and scan-specific model of the hypercapnia achieved by our breath-holding protocol. By determining the voxelwise optimal temporal shift for this model, we produce more accurate estimates of SCVR amplitude and an additional complementary metric of hemodynamic delay. This approach is becoming standard in the field, and BOLD fMRI-based cerebrovascular reactivity studies have increasingly employed a transfer function analysis (gas inhalation) or voxelwise shift optimization procedure (breathing task) to model local hemodynamics ^20–30^.

In this pioneering work, we explore SCVR from two approaches. In the first **(Group SCVR)**, we aim to establish proof-of-concept of our SCVR approach in the cervical spinal cord, mapping the group-level BOLD response from a cohort of 27 healthy participants. Participants completed 2 breath-holding task fMRI runs across 2 sessions. However, to provide more clinically relevant information, SCVR needs to be characterized in individuals. Therefore, in the second study **(Individual SCVR)** we focus on 2 highly-sampled participants to evaluate feasibility of individual SCVR mapping. Each participant completed 18 breath-holding task fMRI runs across 3 MRI sessions.

In both of these approaches, we mapped SCVR and the associated hemodynamic delay using a lagged general linear model (GLM) strategy. To map SCVR amplitude, P_ET_CO_2_ task regressors were fit to the fMRI timeseries in a GLM (also incorporating denoising via nuisance regression). Output parameter estimates (%BOLD per change in P_ET_CO_2_ in mmHg) were combined across the group (Group SCVR) or across individuals’ runs (Individual SCVR). Hemodynamic delay mapping was achieved by temporally shifting the P_ET_CO_2_ regressor and fitting each shifted regressor to the fMRI timeseries. Then, the shift with the best fit across participants (Group SCVR) or runs (Individual SCVR) was used to create a hemodynamic delay map, representing regional variation in the timing of this vascular response across the spinal cord. The SCVR amplitude estimate at this optimal shift was considered the “delay-corrected” SCVR.

We present maps of SCVR amplitude and hemodynamic delay at the group-level and in highly-sampled individuals, in the C5–C8 segments of the human cervical spinal cord. Furthermore, by thresholding the hemodynamic delay maps, we produce noninvasive maps of vascular territories in the human spinal cord, showing agreement with territory maps extrapolated from invasive and preclinical studies. Our scanning methods are readily implemented on most 3T MRI scanners, making this approach to characterizing spinal cord hemodynamics immediately available for basic science and translational research into human spinal cord neurovascular physiology and pathology.

## Results

### Group SCVR amplitude

The breath-hold response was modeled with a P_ET_CO_2_ task regressor in first-level GLMs, allowing SCVR to be mapped in standard semi-quantitative units of %BOLD/mmHg. Group SCVR amplitude was mapped in PAM50 template space **(Fig. 1)**. The group-level SCVR response is distributed predominantly within the ventral spinal hemicord, primarily in or near the gray matter ventral horns, and spread throughout the longitudinal extent of the cervical spinal cord.

**Fig. 1.**
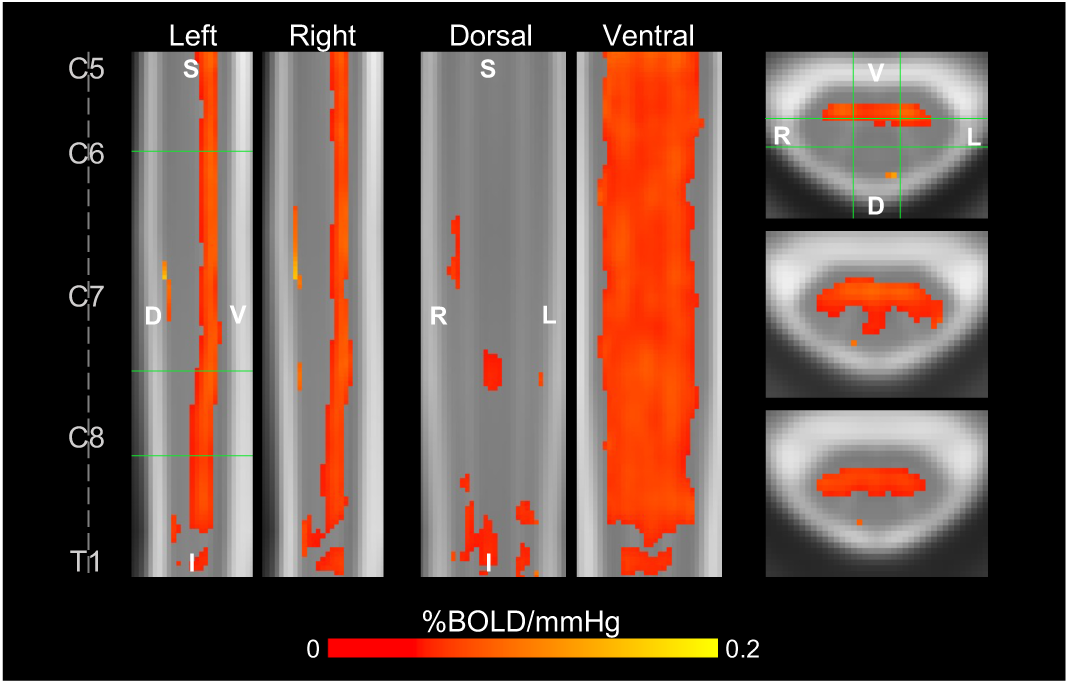
Group-level SCVR amplitude map. Significant SCVR is shown in units of %BOLD/mmHg (*p* < 0.05, FWE-corrected). Two sagittal, two coronal, and three axial slices are shown (green lines indicate location of slices). Approximate centers of spinal cord segments are indicated. (FWE=family-wise error, S=superior, I=inferior, D=dorsal, V=ventral, L=left, R=right).

### Individual SCVR amplitude

The P_ET_CO_2_ regressors show high task compliance across 18 fMRI task runs for the two highly-sampled subjects **(Fig. 2A,E)**. First-level GLMs were calculated, and then the 18 output parameter estimate maps were collated. Individual SCVR amplitude was mapped in PAM50 template space **(Fig. 2B,F)**. SCVR amplitude for both highly-sampled subjects has a similar spatial distribution to the group-level maps, with a high density of significantly responding voxels in the ventral gray matter tissue. However, the amplitude for Subject 1 is visibly higher than for Subject 2.

**Fig. 2.**
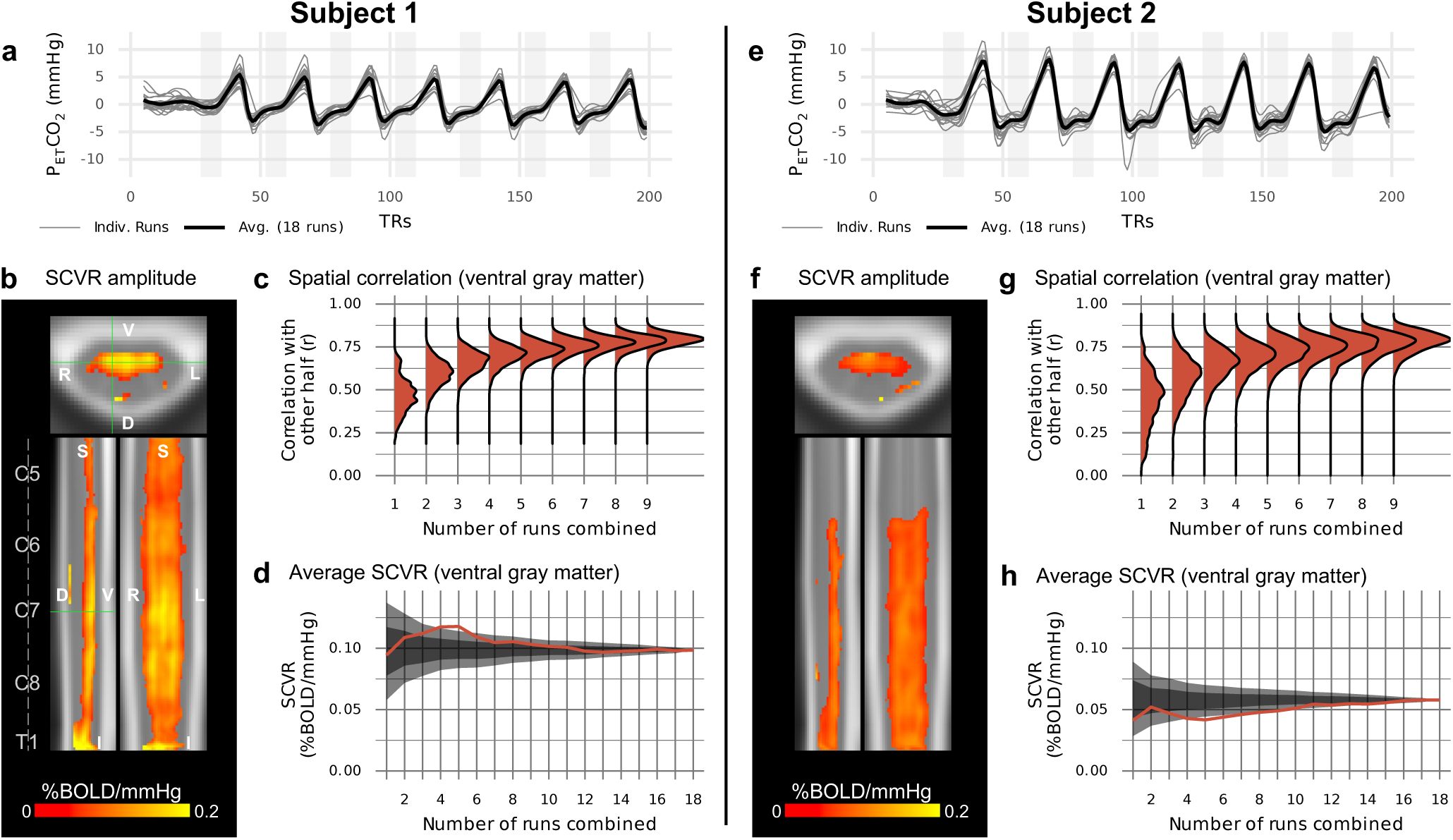
SCVR mapping in two highly-sampled subjects. **(A,E)** P_ET_CO_2_ regressors (convolved with the canonical HRF and demeaned) for 18 breath-holding task runs and their average. **(B,F)** Maps of SCVR amplitude in axial, sagittal, and coronal planes (FWE-corrected, *p* < 0.05). Spinal cord segments and slice locations (green lines) are indicated. **(C,G)** Split-halves spatial correlation of gray matter SCVR amplitude for varying number of runs combined (1-9) vs. other half combined (9 runs). **(D,H)** Average SCVR as runs are sequentially combined in the order they were acquired (red line) and in randomly shuffled orders (shaded areas; ±1SD and ±2SD).

To investigate the reliability of SCVR amplitude estimates generated from subsets of the highly-sampled individual datasets, a split halves analysis was performed. Modeled after Laumann et al. 2015 ^31^, this was done by randomly and repeatedly splitting gray matter SCVR estimates for the 18 runs into two halves and comparing combinations of one half (1-9 runs) to the other half combined (9 runs). A spatial Pearson correlation within ventral gray matter voxels was used to compare the two halves **(Fig. 2C,G)**. In both subjects, comparing SCVR estimates derived from only 1 run versus from 9 runs combined yields moderate spatial correlation (median *r* < 0.5). When comparing SCVR estimates from 6 runs combined versus 9 runs combined, this yields a strong spatial correlation (median *r* ≈ 0.7-0.8). Looking at average SCVR as runs are incrementally combined, the variability initially rapidly decreases, and then more gradually **(Fig. 2D,H)**.

### Group SCVR delay

The distinct ventral distribution of significant SCVR responses raises the question of whether variation in the timing of the SCVR response may be obscuring this effect in more dorsal regions. To investigate and account for temporal delays, first-level models were recalculated with shifted versions of the P_ET_CO_2_ task regressor (±10s, in increments of 2s) **(Fig. 3A)**. Group-level SCVR was calculated for each shift using the same method as above. The optimal shifts were identified, and the corresponding delay map and delay-corrected SCVR amplitude map were generated **(Fig. 3B, Supplementary Fig. S1)**. (Note, this approach assumes that the timing of the voxelwise SCVR response is the same across all subjects.)

**Fig. 3.**
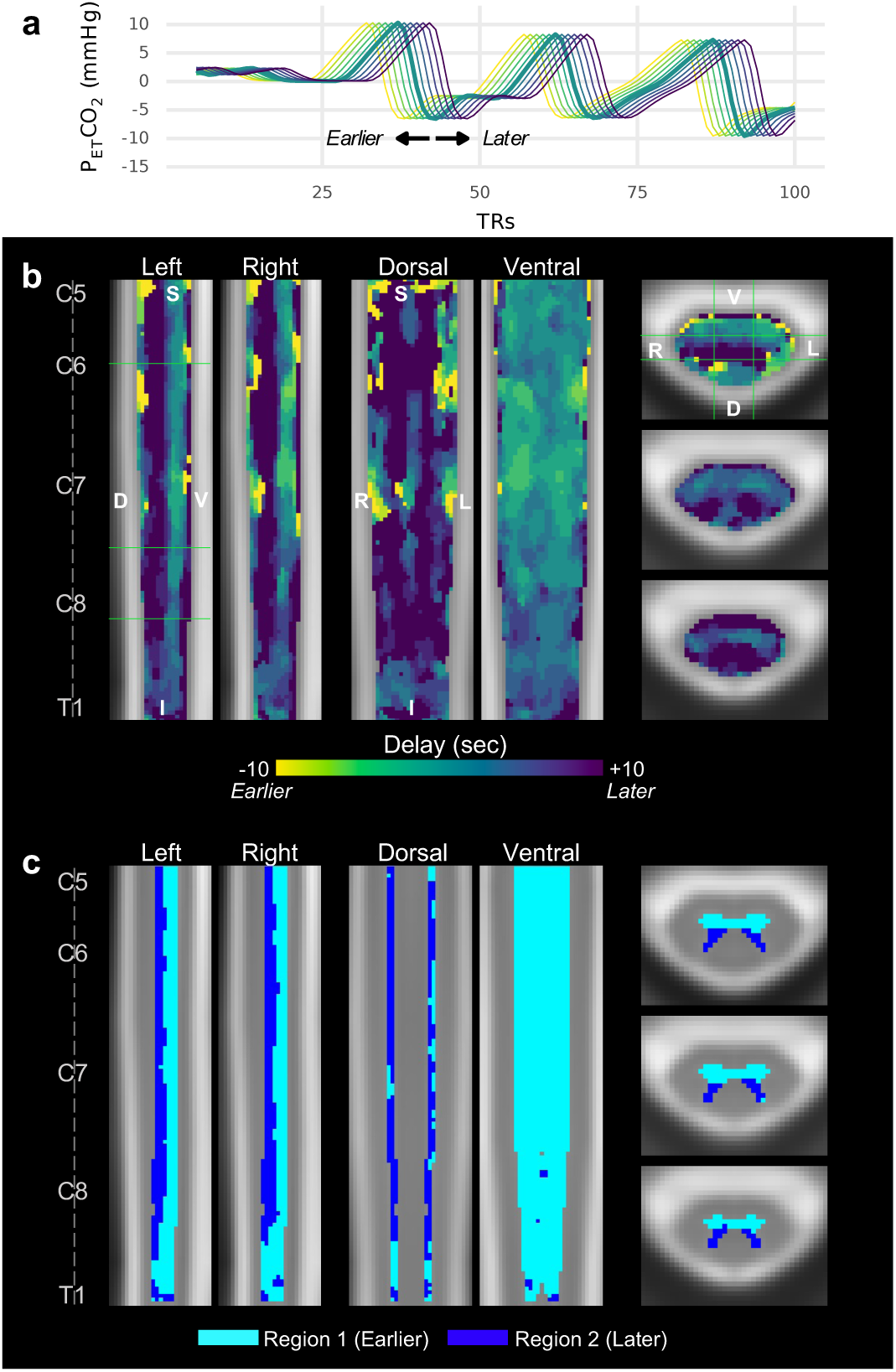
SCVR hemodynamic delay mapping at the group-level. **(A)** Example of temporally shifted P_ET_CO_2_ regressors. The unshifted (no delay) P_ET_CO_2_ regressor (thick green line) and each tested shift ±10 s, in 2s increments, are shown. Line colors correspond to the “Delay” colorbar in panel B. Full traces are truncated at 100 TRs for ease of visualization. **(B)** SCVR hemodynamic delay map (ranging ±10 s, in increments of 2 s). The same slices as Fig. 1 are shown. **(C)** Gray matter regional segmentation based on delay threshold.

As anticipated, the delay-corrected SCVR amplitude map shows a more diffuse distribution of significantly responding voxels. Delay-corrected SCVR (mean ± SD) is 0.04 ± 0.02 %BOLD/mmHg in the cord and 0.05 ± 0.02 %BOLD/mmHg in the ventral gray matter. Across all participants, the distribution of Group SCVR hemodynamic delay shows an earlier response in the ventral cord, and later in the dorsal cord. Moreover, these earlier and later responding regions appear to resemble the shape of the ventral and dorsal gray matter horns, respectively. The mean (± SD) delay between the dorsal and ventral horns is approximately 6.2 ± 4.9 seconds (about 3 times our sampling TR).

Using a histogram of gray matter SCVR delay, a delay threshold was manually defined; each voxel was labeled as Region 1 (sub-threshold, earlier response) or Region 2 (supra-threshold, later response), which comprises the delay-based gray matter segmentation **(Fig. 3C)**. The approximate boundary between the ventral and dorsal horns can be clearly observed.

### Individual SCVR delay

Hemodynamic delay was mapped with the same methods as for Group SCVR. The SCVR delay maps for the two highly-sampled individuals show a similar ventral-dorsal spatial distribution as for Group SCVR **(Fig. 4A,E)**. The delay between dorsal and ventral gray matter is approximately 6.3 ± 5.2 seconds and 7.6 ± 8.0 seconds for Subject 1 and Subject 2, respectively. A delay-corrected SCVR amplitude map was created using the parameter estimates associated with the optimal shift for each voxel **(Fig. 4B,F)**, and shows a more diffuse spread of significantly responding voxels, (compared to Fig. 2B,F), although there are still dorsal regions without a significant response. The mean delay-corrected SCVR in the spinal cord for Subject 1 and Subject 2 is 0.08 ± 0.06 %BOLD/mmHg and 0.07 ± 0.06 %BOLD/mmHg, respectively. And in the ventral gray matter for Subject 1 and Subject 2 is 0.10 ± 0.04 %BOLD/mmHg and 0.07 ± 0.04 %BOLD/mmHg, respectively.

**Fig. 4.**
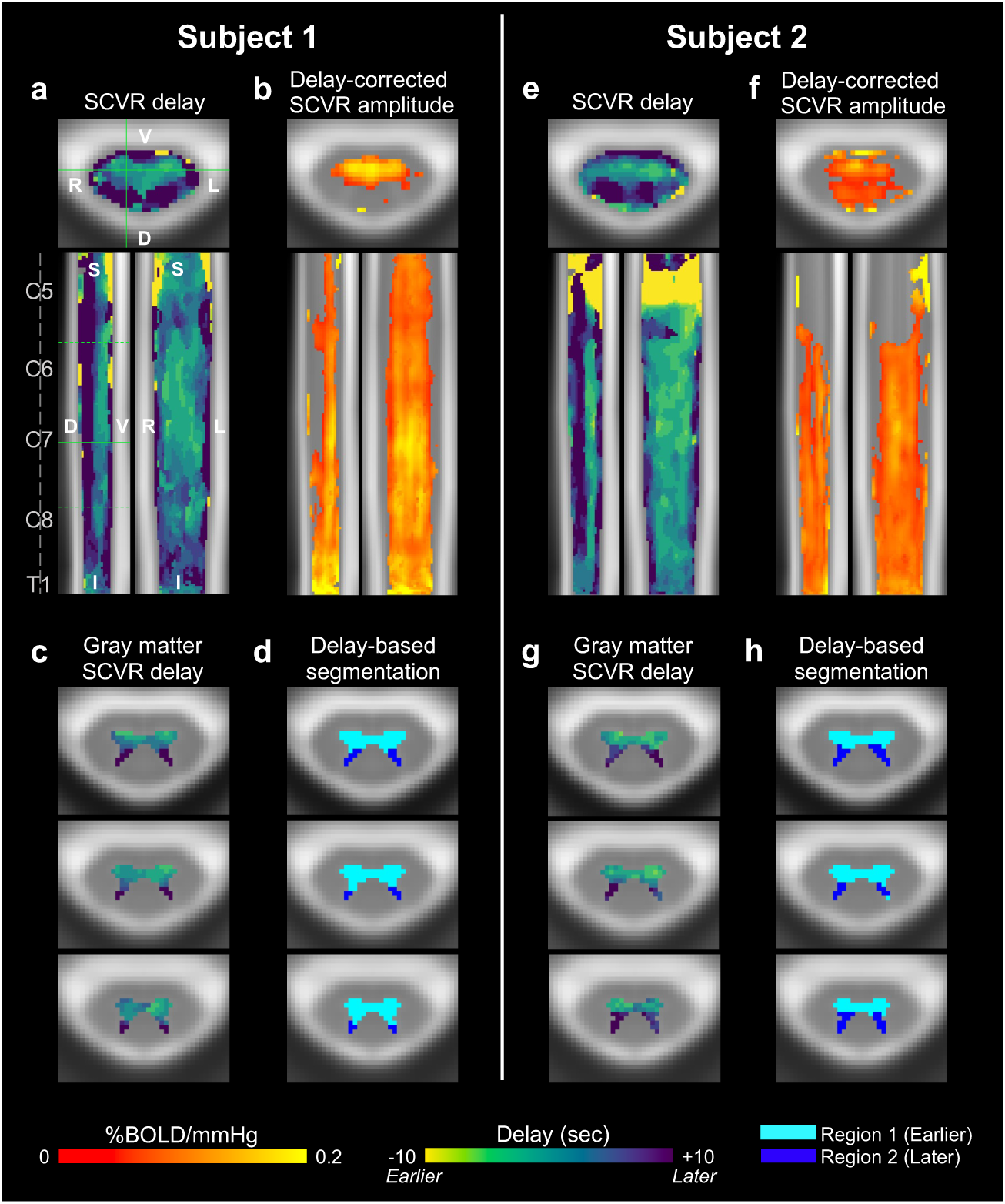
SCVR delay mapping in two highly-sampled subjects. **(A,E)** SCVR delay map (ranging ±10 s, in increments of 2 s). Spinal cord segments and slice locations (solid green lines) are indicated. The three solid and dotted green lines in the sagittal view indicate axial slices in the lower panels. **(B,F)** Delay-corrected SCVR amplitude (FWE-corrected, *p* < 0.05, Šidák corrected). The same slices as Fig. 2 are shown. **(C,G)** SCVR delay map within gray matter voxels. **(D,H)** Gray matter regional segmentation based on delay threshold.

The individual subject SCVR hemodynamic delay maps, restricted to show gray matter voxels, clearly recapitulate the earlier and later responses in the ventral and dorsal horns, respectively **(Fig. 4C,G)**. Applying the thresholding approach described above, the ventral and dorsal boundary is defined, and agrees with the group-level results **(Fig. 4D,H)**.

## Discussion

In this work, we used a breath-holding task to evoke systemic vasodilation and a corresponding BOLD fMRI response in the spinal cord, and then modeled SCVR amplitude and hemodynamic delay. Applying these methods in a cross-sectional cohort and in two highly-sampled individuals, we produced robust spatiotemporal maps of vascular function in the human cervical spinal cord. Our results reveal hemodynamic differences in ventral versus dorsal gray matter regions, consistent across individuals, suggesting that SCVR delay could delineate vascular territories in the cord.

SCVR modeled with P_ET_CO_2_ provides semi-quantitative information about the regional spinal cord hemodynamic response to a vasoactive stimulus. Cerebrovascular reactivity has already proven to deliver clinically relevant information about cerebrovascular health ^16,17^, indicating the potential utility of similar measures for the spinal cord. In the healthy cohort examined in this study, the distribution of significant SCVR (without delay correction) is restricted to more ventral tissue, although extending throughout the longitudinal extent of our imaging field of view. Sensitivity of the SCVR technique in this region may be particularly relevant for early detection of vascular changes in patients with degenerative cervical myelopathy, as age-related cord compression and subsequent neurodegeneration is most prevalent in the ventral regions ^32^.

The significant ventral SCVR response is predominantly, but not strictly, localized to gray matter voxels. This agrees with observations of vascular reactivity in the brain, where the cerebrovascular reactivity amplitude is greater in gray matter versus white matter voxels due to it being more highly vascularized and having higher SNR ^17^.

Early reports of BOLD signal changes in the spinal cord achieved with a controlled hypercapnic gas inhalation challenge were roughly 0.17 %BOLD/mmHg (based on a reported 0.6 ± 0.3 mean percent signal change with a 3.5 ± 0.8 mmHg mean P_ET_CO_2_ change ^18^), as compared to 0.04 ± 0.02 %BOLD/mmHg for the group delay-corrected SCVR across the whole spinal cord region-of-interest in this study. This difference may be due to smaller sample sizes inflating parameter estimates ^33^, or because of the different methods of averaging – our measure is obtained from the group level parameter estimates. Additionally, the response to breath-holding is transient, while gas inhalation evokes a more prolonged response, making delay correction particularly important for breath-holding SCVR. Delay correction performed at the group-level, however, is sub-optimal due to anticipated variability across participants. This is evidenced by the higher mean SCVR estimates for the two highly-sampled subjects. Furthermore, breath-hold induced hypercapnia is associated with a confounding effect of hypoxia, while controlled hypercapnia is not ^34,35^. Based on Schulman and Uludağ’s simulation of cerebrovascular reactivity with and without hypoxia (to mimic breath-holding and controlled hypercapnia, respectively) breath-holding is expected to underestimate cerebrovascular reactivity measurements, which agrees with reports in the literature ^34^. The mean group delay-corrected SCVR in ventral gray matter is 0.05 ± 0.02 %BOLD/mmHg, which is about four to five times lower than similar mean gray matter cerebrovascular reactivity measures ^36,37^ and is to be expected because of higher SNR in the brain.

The SCVR maps derived from the highly-sampled individuals reveal higher SCVR amplitude for Subject 1 compared to Subject 2. There is also a conspicuous absence of significant SCVR estimates in the superior spinal cord for Subject 2. The underlying reason for these differences in SCVR amplitude is unclear and may represent differences in anatomy, physiology, or regional fMRI signal quality.

Individual SCVR amplitude maps represent the 126 breath-holds performed across the 18 fMRI runs. Considering our long-term goal to use SCVR to provide clinical insight into neurological pathology development and progression, it is not reasonable to expect patients to undergo over 100 minutes of breath-hold fMRI. To assess the feasibility of clinical translation using our current SCVR acquisition strategy, we implemented a split halves analysis to probe the number of fMRI runs necessary to achieve reliable SCVR estimates. As shown in **Fig. 2**, the voxelwise spatial correlation in ventral gray matter begins to plateau at approximately 5-6 runs, indicating data from one scan session (34-41 min) may be sufficient to map SCVR. The high initial variability in average ventral gray matter SCVR is also substantially reduced at 5-6 runs combined. Although this scan duration may limit the use of SCVR methodology to research studies, we anticipate that future methodological improvements in acquisition, denoising, and machine learning will further decrease this scan time requirement and make SCVR mapping a viable clinical scan technique. Furthermore, there is relatively low variability in voxelwise SCVR within gray matter in these healthy individuals, and spatial correlations may be much higher in subjects with greater heterogeneity in SCVR values due to pathology.

Since the hypercapnic breath-hold challenge evokes a systemic vasodilatory response, we expected a more widespread fMRI response throughout both ventral and dorsal spinal cord tissue. The absence of significant dorsal responses in **Figs. 1** and **2** allude to spatiotemporal variability in the relationship between our P_ET_CO_2_ model and the associated BOLD response across the spinal cord. Differences in timing could arise because of arterial transit time differences and local variability in the vasodilatory blood flow response and associated BOLD contrast changes. Maps of SCVR hemodynamic delay reveal local regional variation in the timing of the blood flow response that are consistent across our cohort, with earlier and later responses in the ventral and dorsal gray matter, respectively. Although BOLD fMRI is primarily sensitive to venous effects, the response to CO_2_ is driven by arteriolar vasodilation. So, local timing differences may (at least in part) represent territories of arterial supply to the spinal cord.

Supplying approximately the ventral two-thirds of the cord is the <u>central system</u>, derived from the anterior spinal artery (ASA) ^1^. From the ASA, central sulcal arteries penetrate into the anterior median fissure ^1,38^. Blood flows centrifugally, supplying the ventral gray matter, intermediate zone, ventral aspect of the dorsal gray matter, and other white matter regions ^1,39^. This aligns well with the spatial distribution of the earlier ventral gray matter response.

The approximate remaining one third of the cord is supplied by the <u>peripheral system</u>, derived from the paired posterior spinal arteries (PSAs) and pial arterial plexus ^1^. The ASA and PSAs contribute to an anastomosed network on the periphery of the spinal cord, the pial arterial plexus, from which vessels penetrate the spinal cord perpendicular to its surface ^1,39^. Blood flows centripetally, supplying the dorsal aspect of the dorsal gray matter and other white matter regions ^1,39^. This aligns well with the spatial distribution of the later SCVR responses. Additionally, by applying a simple threshold to the SCVR hemodynamic delay maps, we can cleanly delineate these two gray matter vascular territories: (1) areas with an earlier response of likely ASA supply (ventral horns and intermediate zone) and (2) areas with a later response of likely PSAs supply (dorsal horns).

The observed difference in delay between the ventral and dorsal horns was on the order of several seconds for both the group and individual SCVR maps. In interpreting the spatial distribution of delay across the spinal cord, it is important to remember that the SCVR response does not only represent an arterial transit time difference but also represents variation in the local vasodilatory response as well as variation in the BOLD signal evolution. While differences in arterial transit time measured across the brain are about 1-2 seconds ^40,41^, maps of cerebrovascular reactivity delay using BOLD fMRI in healthy participants have shown larger regional differences of several seconds across brain regions ^21,24,27,28^. It is also relevant to consider how these estimates may be influenced by the spatial resolution, particularly in the dorsal horn. Specifically, there is the potential for partial volume effects with the nearby white matter, which may be exacerbated during co-registration and the transformation of images from the subject space to the template space. The capillary bed in white matter is much less dense as compared to the gray matter ^1,42,43^, similar to what is observed in the brain ^17^. Furthermore, a portion of the dorsal horn is not well supplied by the capillary bed ^1^. These factors may adversely affect SCVR amplitude and delay estimation ^17^. Higher magnetic field strengths are an appealing choice for imaging white matter hemodynamic signals ^44^, and future ultra-high field MRI work may offer improved characterization of dorsal horn or white matter SCVR.

These local timing differences in the BOLD response to hypercapnia are driven by multiple sources such as vascular transit delays, variation in the local vasodilatory response, and variation in the BOLD signal response ^20,21,24,27^. Incorporating a voxelwise temporal shift has demonstrated differences in delay across the healthy or injured brain ^21,22,24^ and accounting for these delays can provide for more accurate local estimation of cerebrovascular reactivity, which is particularly relevant for measurement in pathology. Similarly, when accounting for SCVR hemodynamic delay by “correcting” the timing of the P_ET_CO_2_ regressor, we have a more optimal model of vascularly active voxels across the entire cord. After delay correction, significantly responding voxels are more diffusely spread over both ventral and dorsal regions, as compared to the original SCVR amplitude map. This agrees with work in the brain showing increased cerebrovascular reactivity estimates after delay correction ^20^. This effect is more pronounced in the highly-sampled individuals than for the group-level results; this is likely due to inter-individual variation in hemodynamic delay, which makes it sub-optimal to apply the same delay correction to an entire group. However, the limited data available for each subject in the cross-sectional analysis are not sufficient to robustly estimate and correct for their unique voxelwise hemodynamic delay patterns using our current methods.

We demonstrate that our fMRI acquisition and modeling approach can produce consistent and meaningful insight into hemodynamics of the human spinal cord. However, there are numerous challenges in robustly mapping SCVR amplitude and delay that should be considered and addressed in future research.

A breathing task, such as breath-holding, is relatively simple to implement and should cause large, detectable variations in the fMRI signal, but is also likely to cause respiration-induced B_0_ artifacts, time-locked to the task. Fourier-based respiratory RETROICOR ^45^ nuisance regressors were included to aid fMRI denoising. There may be concern that some of the signal of interest would be fitted by respiratory RETROICOR regressors, instead of the task. Ultimately, these regressors were included because physiological noise modeling techniques with RETROICOR have been shown to be beneficial with noisy spinal cord fMRI data ^46^. Additionally, these regressors were included in the first-level model, rather than as a separate denoising regression step. The inclusion of RETROICOR denoising in our study may therefore cause SCVR amplitude to be slightly underestimated, and the impact on delay estimation is not yet clear.

Another consideration for breathing tasks is the potential for motion confounds that are time-locked to respiration ^28,47^. Mitigation of these effects was done through a spinal cord-specific motion correction procedure ^48^ and then by using those output motion traces in the GLM (see Methods). However, we acknowledge that effects of motion may not be entirely accounted for by these procedures. In future work, hypercapnia could be achieved using gas inhalation challenges, which may result in a more controlled hypercapnic stimulus that is not time-locked to ventilation, chest position changes, and subsequent B_0_ effects. Gas inhalation challenges could also increase task compliance, because participants do not need to actively comply with the voluntary breathing task. Indeed, the early spinal cord fMRI work involving a hypercapnic stimulus used different gas inhalation techniques ^18,19^. However, the requisite facemask may be less comfortable for participants and less compatible with some coils (e.g., the 64-channel head/neck coil used in this study). Although literature from cerebrovascular reactivity mapping in the brain suggests that these two methods generate similar results ^49^, future work should be done to compare breath-holding task-based SCVR measurements with those derived from gas inhalation challenges.

The ground truth of our SCVR measurements across spinal cord vascular territories is unknown; it will be important in future work to refine the range and step size of the shifted P_ET_CO_2_ regressors. The ±10s P_ET_CO_2_ range was informed by previous work from our lab mapping cerebrovascular reactivity using a shift range of ±9s ^27,50^. It is pertinent to note that histograms of gray matter delay showed a high density at the upper boundary of the range (+10s) as well as the presence of voxels with −10s delays directly adjacent to voxels with +10s delays. These voxels may not reflect physiologically plausible SCVR delays. Additionally, in brain literature, with finer (∼0.3s) increments, shifts at the upper and lower boundary of the range are sometimes not included because they are not considered optimized ^20,27^. Considering the coarse 2s shift increment used in this study, boundary delays were not removed. Refinement of the range and step size will be an important target for future work, and evidence from brain literature suggests this may be particularly important in individuals with pathologically delayed blood supply to the cord ^27^. Additionally, Bayesian modeling of SCVR amplitude and delay, using our initial work to inform spatial priors, could also improve our ability to robustly map individual subject hemodynamics ^25^.

Finally, the P_ET_CO_2_ regressors used throughout this study were convolved with the two-gamma variate canonical HRF ^51^. While this HRF was designed to convert a neural task design into a physiological response, the canonical HRF convolved with P_ET_CO_2_ is often used throughout the cerebrovascular reactivity literature ^16,27,36,52^. In a direct comparison of a set of models, P_ET_CO_2_ convolved with the canonical HRF plus its temporal derivative was shown to be the best model of the BOLD response to breath-holding ^53^. While the present study did not incorporate a temporal derivative, temporal offsets in the local BOLD response were instead accounted for by the analysis of hemodynamic delay. Aside from the delay, there may still be differences in the shape of the response that should be accounted for. In fact, a recent preprint comparing models with voxelwise lag optimization suggests either non-convolved P_ET_CO_2_ or single gamma HRF-convolved P_ET_CO_2_ as the best model. Given the temporal resolution and low SNR of the spinal cord fMRI data, it is not possible to estimate the response shape in this initial study. Future work should explore an appropriate response function specifically for measures of vascular reactivity in the spinal cord.

## Conclusion

SCVR amplitude and delay mapping has the potential to provide clinically relevant information about spinal cord vascular health. Robust maps of SCVR amplitude and delay were presented at the group-level and in two highly-sampled individuals. Hemodynamic delay maps represent the temporal difference in the BOLD response across the cord and resemble expected territories of arterial supply. Future work to refine these promising noninvasive methods will facilitate our long-term goal to establish SCVR as a clinically relevant metric of spinal cord vascular health.

## Methods

### Participants

This study was approved by the Northwestern University Institutional Review Board, and informed consent was obtained for all participants; all methods were performed in accordance with relevant guidelines and regulations. Two datasets were collected for this study and will be referred to as Group SCVR and Individual SCVR. <u>Group SCVR dataset</u>: MRI data were collected from 30 healthy participants (26±5y, 19M). Data were excluded for 3 participants due to excessive image artifacts (N=2) and an incidental finding (N=1). All subsequent Group SCVR analyses represent data from the remaining 27 participants (26±5y, 18F). <u>Individual SCVR dataset</u>: MRI data were collected from 3 healthy participants (24±3y, 2F). Data were excluded for one participant due to radiofrequency image artifacts. Subsequent Individual SCVR analyses represent data from the remaining 2 participants (26±2y, 2F).

### Data Collection

Spinal MRI data were acquired on a 3T Siemens Prisma MRI system (Siemens Healthcare, Erlangen, Germany) with a 64-channel head/neck coil. A SatPad^TM^ cervical collar (SatPad Clinical Imaging Solutions, West Chester, PA, USA) was positioned around the posterior of the neck to increase the homogeneity of the magnetic field around the imaging region. <u>Group SCVR</u>: MRI was acquired before and after the administration of a 30-minute acute intermittent hypoxia (AIH) protocol, as part of a larger research study. <u>Individual SCVR</u>: Anatomical and functional scans were acquired in 3 sessions across 2 days during the same time of day for each participant. Sessions 1 and 2 took place on the first scan day and were in the morning and afternoon, respectively; session 3 took place in the morning on the second scan day. The two scan days were one or two days apart.

### Acute Intermittent Hypoxia (AIH)

AIH has been effective at improving motor function in spinal cord injured cohorts ^54,55^. The data collected for the Group SCVR study was a part of a broader study to investigate with MRI the mechanisms of neural plasticity associated with AIH. This AIH protocol was administered after the first MRI session with a HYP 123 oxygen generator (Hypoxico, Inc., New York, NY, USA) in 15 2-minute cycles. Each cycle consisted of brief exposures to a hypoxic gas mixture (9% FiO_2_, 30-60 seconds), alternating with normal room air (21% FiO_2_, 60-90 seconds), targeting 85% SpO_2_ during each bout. The second MRI session occurred 45-60 minutes after AIH. Note, AIH had a non-significant impact on SCVR (via a two-tailed paired t-test using threshold-free cluster enhancement (TFCE) and 5000 permutations (*randomise*) ^56–58^. Pre- and post-AIH MRI scans were also considered independently to further probe any possible impact of AIH. For details of this analysis see Supplementary Information Section 1. The SCVR maps and average values were consistent (**Supplementary Fig. S2, S3, and S4**). Thus, there was no evidence that AIH modulates SCVR in healthy individuals and the impact of AIH was considered negligible in further analyses.

### Imaging Protocol

A high resolution anatomical T2-weighted scan, covering the brainstem to upper thoracic spine, was acquired with the following parameters: repetition time (TR)/echo time (TE) = 1500/135 ms, sagittal slice thickness = 0.8 mm, in-plane resolution 0.39 mm^2^, 64 slices, flip-angle (FA) = 140°, field-of-view (FOV) = 640 mm^2^. Spinal cord fMRI scans were collected using a gradient-echo echo-planar imaging (EPI) sequence and ZOOMit selective excitation (TR/TE = 2000/30 ms, axial in-plane resolution = 1 mm^2^, axial slice thickness = 3 mm, 25 ascending interleaved slices, FA = 90°, FOV = 128 x 44 mm^2^). The ZOOMit acquisition reduces the field-of-view around the spinal cord. The functional acquisition volume was positioned perpendicular to the spinal cord; the bottom of the volume was positioned at the bottom of the C7 vertebral level. Cervical coverage was approximately from the C4 to C7 vertebral level, but the exact coverage varied due to participant height and spinal anatomy. For each scan session, the task paradigm was displayed to participants via a mirror on the head coil, reflecting a monitor placed behind the bore of the magnet. <u>Group</u> <u>SCVR</u>: A spinal cord anatomical and breath-holding task functional scan were acquired during each of the 2 MRI sessions. Additional spinal cord and brain scans were also acquired during the sessions and were not analyzed in this study. <u>Individual SCVR</u>: A spinal cord anatomical and 5-7 breath-holding task functional scans were acquired during each of the 3 MRI sessions. (It was intended that 6 runs would be collected during each session but due to time constraints during one session, five runs were collected during session 1 and 7 runs were collected during session 2 for Subject 1.) The total number of task runs was 18 for each participant.

### Breath-holding Task Paradigm

A hypercapnic breath-holding task was used to modulate participants CO_2_. The fMRI acquisition was 6 minutes and 50 seconds (205 volumes). The task began with a 20 second rest, followed by 7 breath-hold cycles, and ended with a 30 second rest period. Each breath-hold cycle was as follows: 24 seconds paced breathing (3 seconds in, 3 seconds out), 18 second end-expiration breath-hold, 2 second exhalation, 6 second recovery.

### Physiological Monitoring

Physiological data were collected throughout the fMRI scans, including exhaled CO_2_ and O_2_ through a nasal cannula and breathing via respiratory belt. A pulse transducer was placed either on the dorsalis pedis artery of the foot (for the Group SCVR study because participants were holding hand-grips for another task) or on the right index finger (for the Individual SCVR study). These signals were fed through a Gas Analyzer (CO_2_, O_2_ only) and PowerLab and recorded with LabChart (ADInstruments, Dunedin, New Zealand). The scanner trigger was also recorded through the same system to facilitate alignment of all recordings with fMRI timeseries data.

### fMRI Preprocessing Pipeline

Images were converted from DICOM to NIFTI format (dcm2niix_afni ^59^). <u>Motion Correction:</u> 2D slicewise motion correction was applied with the Neptune Toolbox ^48^ (version 1.211227) and three steps of the toolbox were used to complete motion correction, including the manual definition of a “not cord” mask around the spinal cord and cerebrospinal fluid (CSF) region (#5), computation of a Gaussian weighting mask of the spinal cord, derived from the “not cord” mask (#7), and the application of motion correction (#8). The motion correction algorithm used the Gaussian mask as a weight for each voxel, a temporal median image as the target image for the correction algorithm, and used AFNI (version 22.0.05) to compute and apply the motion parameters to the data (*3dWarpDrive*, *3dAllineate*) ^60,61^. Temporal median filtering was de-selected. Motion corrected functional data and slicewise X and Y motion traces were output. <u>Registration:</u> Co-registration between native, anatomical, and PAM50 template space ^62^ was performed with the Spinal Cord Toolbox (version 5.3.0) ^63^. (Note, the Spinal Cord Toolbox version 6.1.0 included an update to the spinal cord segments and this new version was used to add correct labels to figures.) A binary spinal cord mask in the functional image space was manually drawn (excluding the top/bottom slices) and is used in several steps. The anatomical T2-weighted image of each subject was segmented (*sct_deepseg_sc*) ^64^ and then registered to PAM50 template space (*sct_register_to_template*). The binary spinal cord mask in functional image space and anatomical-template warping field were used to inform registration of the motion corrected functional data to the PAM50 template space (*sct_register_multimodal*). These warping fields were calculated but not applied until after subject-level modeling. <u>Smoothing</u>: Functional data were smoothed using a “restricted smoothing” technique ^65^, within a mask of the spinal cord. A 2 x 2 x 6 mm^3^ full-width half-maximum (FWHM) Gaussian smoothing kernel was used (*3dBlurInMask*). <u>Trimming fMRI Volumes</u>: The first and last 5 volumes were trimmed from the functional data to allow for shifts of the task regressor for hemodynamic delay mapping (discussed later).

### Task and Nuisance Regressors

Physiological data (CO_2_, pulse, and respiratory belt) were preprocessed in a bespoke MATLAB (MathWorks, Natick, MA, R2019b) script. The results of a peak-finding algorithm were manually verified. <u>Breath-holding Task</u>: From the preprocessed CO_2_ data, an end-tidal CO_2_ (P_ET_CO_2_) task regressor was calculated and convolved with the two-gamma variate canonical hemodynamic response function (HRF) ^51^. <u>Respiratory and Cardiac</u>: From the preprocessed respiration belt and pulse transducer data, 8 respiratory and 8 cardiac slicewise RETROICOR ^45,46^ regressors were calculated. Only cardiac regressors were calculated for one subject in the Group SCVR study without belt data. <u>CSF</u>: An initial CSF mask was defined; high variance voxels (top 20% in each slice) were retained in the mask and the average timeseries within this mask was used to create slicewise CSF regressors ^46,66^.

### fMRI Analysis and Statistics

<u>First-level Analysis</u>: FSL ^67^ (version 6.0.3) was used to calculate the first-level fMRI models (FEAT ^68^). The first-level models contained the P_ET_CO_2_ task regressor and 19 nuisance regressors (2 motion, 16 RETROICOR, 1 CSF). All input regressors were demeaned. In FEAT, a lenient high-pass filter cutoff of 100 seconds to chosen to remove the low frequency noise and FMRIB’s Improved Linear Model (FILM) prewhitening was used. One statistical contrast was defined: P_ET_CO_2_>0. The output “contrast of parameter estimate” (COPE) maps were normalized by the voxelwise mean to convert to standard units (i.e., %BOLD/mmHg). Output files were warped from subject functional space to the PAM50 template space. <u>Higher-level Analysis</u>: The COPE maps were averaged across the two sessions within each subject (Group SCVR). A group spinal cord mask was calculated as the consensus region of the co-registered first-level results, and higher-level analyses were performed within this mask, only. This mask was calculated separately for the Group SCVR and Individual SCVR studies. A non-parametric one-sample t-test using TFCE and 5000 permutations was used to calculate the higher-level activation maps with family-wise error (FWE) rate correction (*randomise*) ^56–58^. The average parameter estimates are the SCVR amplitude.

### Reliability of Individual SCVR Estimates

To probe the reliability of SCVR estimates for different amounts of data, a split-halves analysis was used ^31^. For each subject, the 18 COPE maps were randomly split into 2 halves, comparing subsets of one half (1-9 runs) to the other 9 runs combined via spatial Pearson correlation in the ventral gray matter. This was repeated 1000 times to build a distribution. Average SCVR in the ventral gray matter was also considered by incrementally combining runs from 1 run to all 18 runs. This was done in the originally acquired order of fMRI runs and then for 100 randomly shuffled orders.

### SCVR Hemodynamic Delay

10 temporally shifted versions of the P_ET_CO_2_ regressor were created to allow for timing differences of ±10 s in the increment of the TR (2 s). First-level models were repeated identically to the original except for the temporally shifted P_ET_CO_2_ regressor. The higher-level analyses in PAM50 space were also repeated for each shift. For each voxel, the highest t-statistic across the delay range was designated as the optimal shift (the “delay” for that voxel) (*getBestFits)*. <u>Delay-corrected SCVR Amplitude</u>: These optimal shifts were then used to create a delay-corrected SCVR map. For each run, delay-corrected COPE maps were generated using the parameter estimates associated with the optimal shift (*delayCorrectedSCVR*). This resulted in delay-corrected COPE maps for each run, which were then processed with the same higher-level analyses used previously. The average parameter estimates are the delay-corrected SCVR amplitude. Šidák correction was used to account for the multiple tests in the delay-correction procedure ^69,70^. <u>Delay-based</u> <u>Segmentation</u>: A histogram of gray matter delays was manually thresholded. Voxels with delays on either side of the threshold were designated as *Region 1* (Earlier) or *Region 2* (Later).

## Acknowledgements

This work was supported by the Center for Translational Imaging at Northwestern University. The authors would like to thank Robert L. Barry for contributions to the development of our data preprocessing pipeline and Sameer Faruquee for contributions to data preprocessing. Research supported by the Craig H. Neilsen Foundation (595499). KJH was supported by an NIH NIBIB-funded training program (T32EB025766) and NINDS-funded predoctoral fellowship (F31NS134222). This research was supported in part through the computational resources and staff contributions provided for the Quest high performance computing facility at Northwestern University which is jointly supported by the Office of the Provost, the Office for Research, and Northwestern University Information Technology.

## Author Contributions

M.G.B., M.S.S., and T.B.P. designed research; K.J.H. and M.A.H. performed research; K.J.H. analyzed data; K.J.H. wrote the initial draft; all authors reviewed the manuscript.

## Code availability statement

The code to perform the SCVR delay analysis is available at https://github.com/BrightLab-ANVIL/SCVR.

## Additional Information

The authors declare no competing interests.

