## Supplementary Material for "MRI mapping of hemodynamics in the human spinal cord"

### Supplementary Information

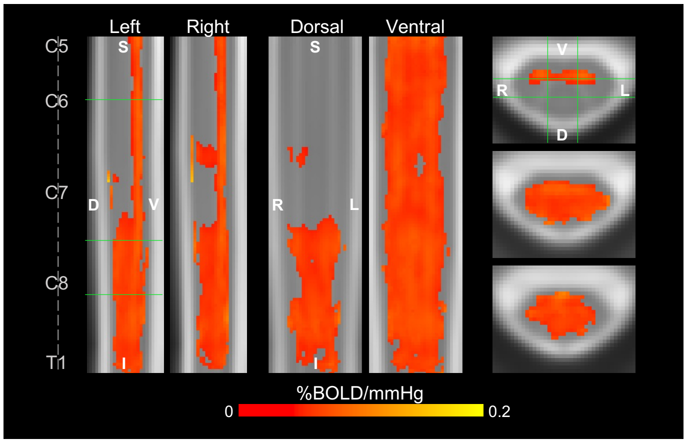


**Supplementary Fig. S1. Group SCVR delay-corrected amplitude map.** Delay-corrected SCVR amplitude (family-wise error (FWE) rate-corrected, $p < 0.05$, Šidák corrected). Axial, sagittal, and coronal slices are the same as those shown in Fig. 1. (S=superior, I=inferior, D=dorsal, V=ventral, L=left, R=right)

### 1 SCVR and Acute Intermittent Hypoxia (AIH)

#### *1.1 Voxelwise t-test for AIH*

Although AIH has been shown to be effective in improving motor function after spinal cord injury [1,2], no significant effect was detected when comparing pre-AIH (Scan 1) vs. post-AIH (Scan 2) SCVR maps. Specifically, no significant voxels were identified using a non-parametric two-tailed paired *t*-test with 5000 permutations and threshold-free cluster enhancement (TFCE) (*randomise*). (Similarly, no significant effect of AIH was found in our previous work characterizing spinal cord hand-grasping motor function in healthy individuals[3].)

#### *1.2 Mapping SCVR Pre- and Post-AIH*

To further probe any possible impact of AIH the pre- and post-AIH scans were also considered independently and SCVR maps were generated. This analysis was nearly identical to the described statistical analyses in the Methods (*fMRI Analysis and Statistics* section). Briefly, pre-AIH or post-AIH first-level fMRI COPE maps were collated across subjects in PAM50 template space. A non-parametric one-sample t-test using TFCE and 5000 permutations was used to calculate group-level maps within a group spinal cord mask. SCVR hemodynamic delay and delay-corrected amplitude maps were generated as described in the Methods (*SCVR Hemodynamic Delay* section).

**Supplementary Figure S2** shows the group-level SCVR amplitude maps for each the pre-AIH and post-AIH scans. Notably, the strong ventral response observed first in **Fig. 1** is still present in both of these maps. The spatial distribution of significant voxels and the magnitude of SCVR estimates appears overall similar across the maps presented in **Fig. 1** and **Fig. S2**. Similarly, the group SCVR delay map **(Fig. S3)** and delay-corrected SCVR amplitude map **(Fig. S4)** both have similar spatial distributions compared to the original delay map **(Fig. 3B)** and delay-corrected map **(Fig. S1)**, respectively.

The average SCVR estimates were also consistent between scans. Specifically, the whole cord mean delay-corrected SCVR for the pre- and post-AIH scans are 0.041 ± 0.028 %BOLD/mmHg and 0.036 ± 0.026 %BOLD/mmHg, respectively. The ventral gray matter mean delay-corrected SCVR for the pre- and post-AIH scans are 0.046 ± 0.020 %BOLD/mmHg and 0.047 ± 0.019 %BOLD/mmHg, respectively. Therefore, no evidence was found that AIH affected SCVR estimates in these healthy individuals.

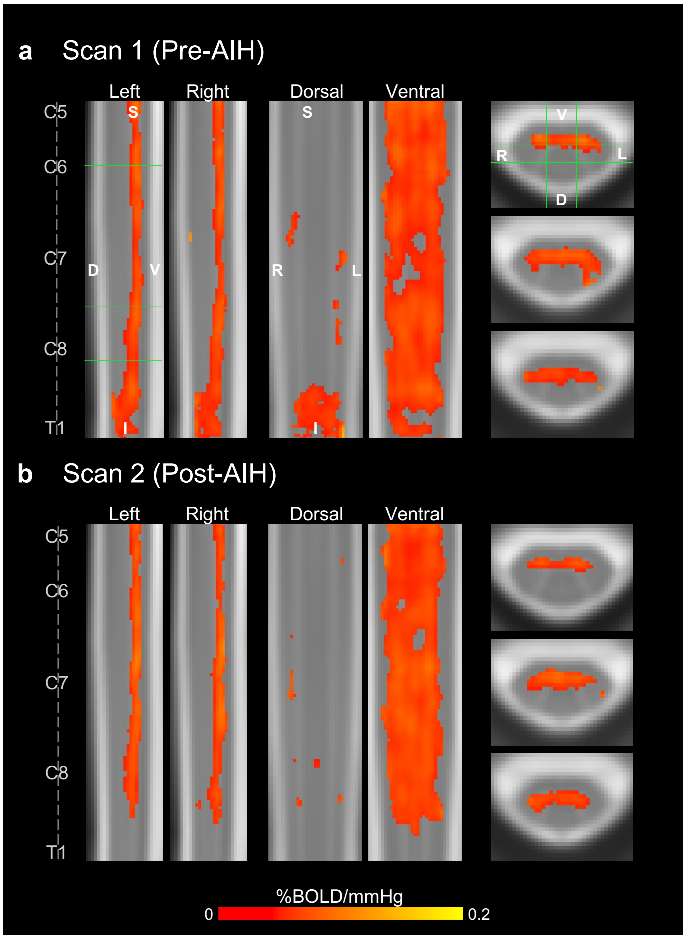


**Supplementary Fig. S2. Group-level SCVR amplitude maps for Pre- and Post-AIH scans.** Significant SCVR is shown in units of %BOLD/mmHg ($p$ < 0.05, FWE-corrected). Axial, sagittal, and coronal slices are the same as those shown in Fig. 1.

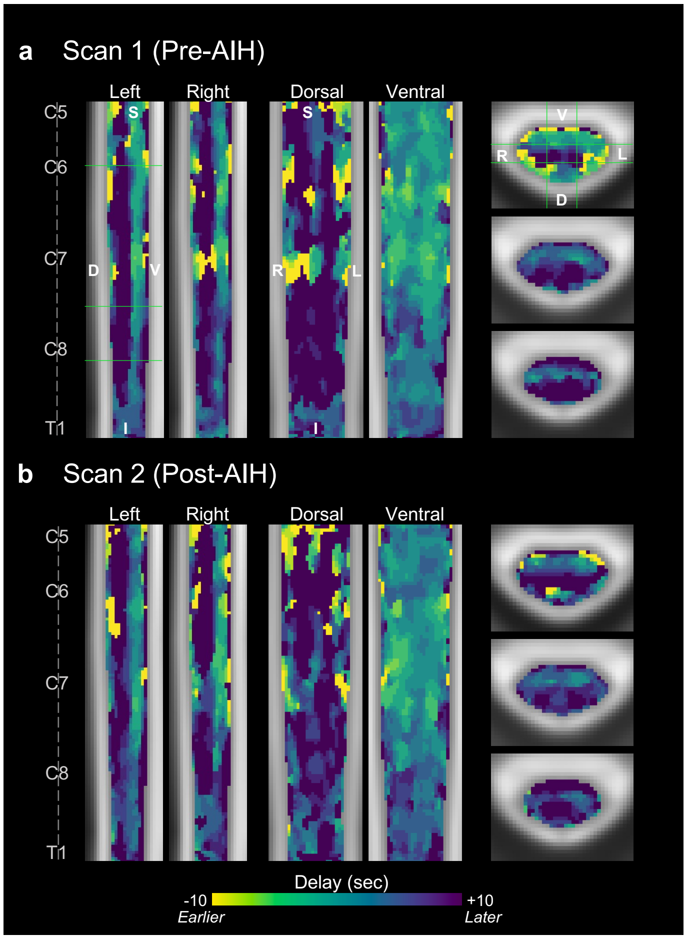


**Supplementary Fig. S3. Group-level SCVR hemodynamic delay maps for Pre- and Post-AIH scans.** Delays range ±10 s, in increments of 2 s. Axial, sagittal, and coronal slices are the same as those shown in Fig. 1.

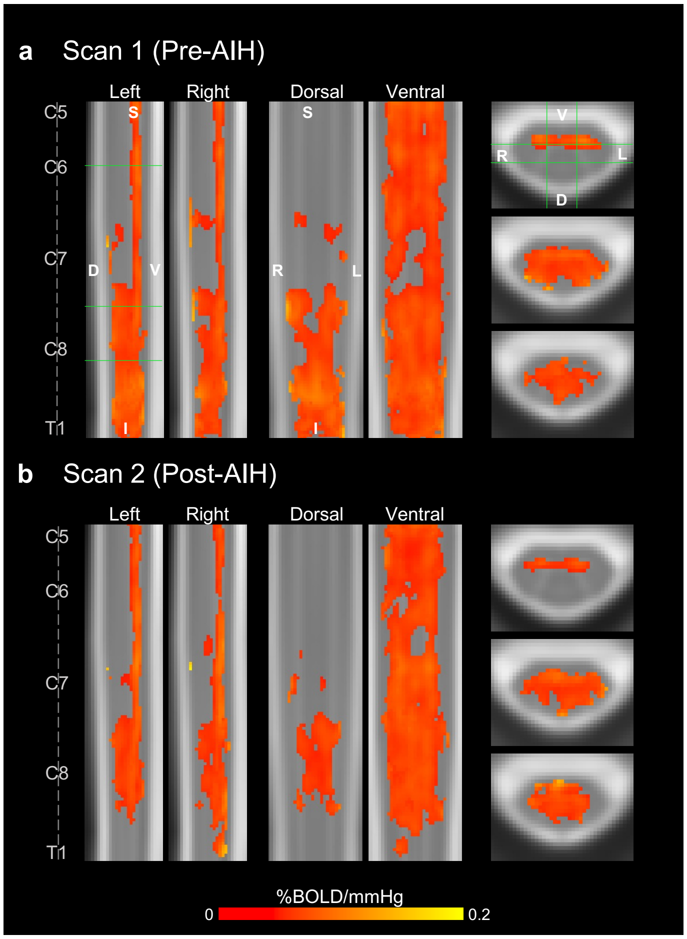


**Supplementary Fig. S4. Group-level SCVR amplitude maps for Pre- and Post-AIH scans.** Significant SCVR is shown in units of %BOLD/mmHg ($p < 0.05$, FWE-corrected, Šidák corrected). Axial, sagittal, and coronal slices are the same as those shown in Fig. 1.
